# High-fat diet intake modulates maternal intestinal adaptations to pregnancy, and results in placental hypoxia and impaired fetal gut development

**DOI:** 10.1101/436816

**Authors:** Wajiha Gohir, Katherine M. Kennedy, Jessica G. Wallace, Michelle Saoi, Christian J. Bellissimo, Philip Britz-McKibbin, Jim J. Petrik, Michael G. Surette, Deborah M. Sloboda

## Abstract

Shifts in maternal intestinal microbiota have been implicated in metabolic adaptations to pregnancy. In this study we investigated how high-fat diet intake impacts the maternal gut microbiota, intestinal inflammation and gut barrier integrity, placental inflammation, and fetal intestinal development at E18.5. High-fat diet (HFD) was associated with decreased relative abundancesof SCFA producing genera during pregnancy. These diet-induced shifts paralleled decreased maternal intestinal mRNA levels of SCFA receptor *Gpr41*, modestly decreased cecal butyrate, and altered mRNA levels of inflammatory cytokines and immune cell markers in the maternal intestine. Maternal HFD resulted in impaired gut barrier integrity, with corresponding increases in circulating maternal levels of LPS and TNF.Placentafromhigh-fat fed damsdemonstrated blood vessel immatu-rityand hypoxia, decreased freecarnitine, acylcarnitine derivatives, TMAO, as well as altered mRNA levels of inflammation, autophagy and ER stress markers. HFD exposed fetuses had increased activation of NF-κB and inhibition of the unfolded protein response in the developing intestine. Together, these data suggest that high-fat diet intake prior to and during pregnancy shifts the composition of the maternal gut microbiota and impairs gut barrier integrity, resulting in increased maternal circulating LPS, which may ultimate contribute to changes in placental vascularization and fetal gut development.

**Funding information:** Farncombe Family Digestive Health Research Institute (KMK); Canadian Institute of Health Research (CJB); Canada Research Chairs Program (MGS, DMS); Natural Sciences and Engineering Research Council of Canada, Genome Canada (PBM).

## 1 INTRODUCTION

Maternal obesity and excess gestational weight gain are key predictorsofchildhood obesity and metabolic complications in adulthood. The link between maternal and offspring obesity has been extensively investigated in both clinical and experimental settings (Gluckman and Hanson, 2007). Excess intake of saturated fats and obesity are associated with increased risk of pregnancy complications including preeclampsia (Triunfo and Lanzone, 2014), and may result in fetal overgrowth and altered placental development (Wallace et al., 2012). Although animal studies have shown maternal high-fat diet (HFD) induced obesity is strongly associated with offspring metabolic disease (Morris and Chen, 2009; Howie et al., 2009; Elahi et al., 2009), the signaling pathways by which maternal HFD intake, obesity,or excess gestational weight gain can confer offspring metabolic dysfunction are still unclear.

The relationship between intestinal microbes and host metabolism has become one of most studied factors mediating obesity risk (Holmes et al., 2012), and it has been shown that pregnancy shifts the abundance and type of bacteria that colonize the intestine (Koren et al., 2012). These shifts have been suggested to contribute to maternal metabolic adaptations to pregnancy and may mediate pregnancy outcomes (Koren et al., 2012; Goltsman et al., 2018). We, and others, have shown that these bacterial shifts are further modified by maternal HFD and obesity (Collado et al., 2008; Santacruz et al., 2010; Gohir et al., 2015), but the signaling molecules that relate microbial abundance to inflammatory and metabolic responses during pregnancy have not been thoroughly investigated. Recent data show that maternal propionate is negatively associated with maternal leptin and measures of infant weight (Priyadarshini et al., 2014) and suggest that shifts in the pregnant gut microbiota could influence metabolism via alterations in short-chain fatty acid (SCFA) production. SCFAs influence gut barrier function (Burger-van Paassen et al., 2009). Changes in the gut microbiota of male high-fat fed mice are associated with increased circulating bacterial lipopolysaccharide (LPS), inducing metabolic endotoxemia (Cani et al., 2008). In the context of pregnancy, maternal metabolic endotoxemia may contribute to placental inflammation as a result of maternal HFD and obesity (Li et al., 2013; Pantham et al., 2015; Salati et al., 2018).

In this study we aimed to determine the impact of HFD intake prior to and during pregnancy on the maternal intestinal microbiota and its relationship with intestinal inflammation and barrier integrity, and further investigate whether maternal endotoxemia is associated with placental metabolic stress and inflammation, and impaired fetal gut development.

## 2 METHODS

### 2.1 Ethics approval

All animal procedures for this study were approved by the McMaster University Animal Research Ethics Board (Animal Utilization Protocol 12-10-38) in accordance with the guidelines of the Canadian Council of Animal Care and the ethical principles of this journal (Grundy, 2015).

### 2.2 Animal model

Following 1 week of acclimation to the McMaster Central Animal Facility, 4-week-old female C57BL/6J mice (RRID: IMSR_JAX:000664) were randomly assigned to a control (n=10; 17% kcal fat, 29% kcal protein, 54% kcal carbohydrate, 3.40 kcal/g; Harlan 8640 Teklad 22/5 Rodent Diet)or HFD (n=10; 60% kcal fat, 20% kcal protein, 20% kcal carbohydrates, 5.24 kcal/g, Research Diets Inc. D12492). Females were housed two per cage with ad libitum food and water at a constant temperature (25°C) on a 12:12 light-dark cycle. Following 6 weeks of dietary intervention, fecal samples were collected from each mouse (identified as before mating; BM) and were co-housed with a control-fed C57BL/6J male. Mating was confirmed by visualization of a vaginal plug and designated embryonic day (E)0.5. Pregnant females were housed individually, and gestational weight and food intake were recorded throughout pregnancy. Maternal fecal samples were collected at embryonic day (E)0.5, E10.5, E15.5 and E18.5. On E18.5, maternal serum was collected via tail vein blood, and stored at −80°C for future analyses. Pregnant mice were sacrificed by cervical dislocation. Maternal cecal contents and intestinal tissue from the duodenum, jejunum, ileum, and colon snap-frozen in liquid nitrogen and stored at −80°C. Fetuses and placental tissues were dissected, sexes were identified, and fetal weights were recorded. Fetal small and large intestines were snap-frozen in liquid nitrogen and stored at −80°C. Placentas from each litter were snap-frozen in liquid nitrogen and fixed in 4% paraformaldehyde. All sample sizes in the study represent the mother (dam) as a biological replicate.

### 2.3 Maternal microbiota profiling

#### DNA extraction and 16S rRNA gene sequencing

Genomic DNA was extracted from fecal samples as previously described (Whelan et al., 2014) with some modifications: 0.2 g of fecal material was used and additional mechanical lysis step was included using 0.2 g of 2.8 mm ceramic beads. PCRamplificationofthe variable3(V3)region of the 16S rRNA gene was subsequently performed on the extracted DNA from each sample independently using methods described previously (Bartram et al., 2011; Whelan et al., 2014). Each reaction contained 5pmol of primer, 200 mmol of dNTPs, 1.5μl50mM MgCl_2_,2μl of 10 mg/ml bovine serum albumin (irradiated with a transilluminator to eliminate contaminating DNA) and 0.25μl Taq polymerase (Life Technologies, Canada) for a total reaction volume of 50μl. 341F and 518R primers were modified to include adapter sequences specific to the Illumina technology and 6-base pair barcodes were used to allow multiplexing of samples as described previously. 16S DNA products of the PCR amplification were subsequently sequenced using the Illumina MiSeq platform (2×150bp) at the Farncombe Genomics Facility (McMaster University, Hamilton ON, Canada).

#### Processing and analysis of sequencing

Resultant FASTQ files were processed using a custom in-house pipeline as previously described (Whelan et al., 2014). Briefly, reads exceeding the length of the 16S rRNA V3 region were trimmed (cutadapt, RRID:SCR_011841), paired-end reads were aligned (PANDAseq, RRID:SCR_002705), Operational Taxonomic Units (OTUs) were grouped based on 97% similarity (AbundantOTU+, SCR_016527). Taxonomy was assigned the RDP Classifier (Ribosomal Database Project, RRID:SCR_006633) against the Feb 4, 2011 release of the Greengenes reference database (Greengenes, RRID:SCR_002830). Any OTU not assigned to the bacterial domain were culled, as was any OTU to which only 1 sequence was assigned. This processing resulted in a total of 11,027,452 reads (mean 137,843 reads per sample; range: 59,233-249,890) and 9237 OTUs.

Taxonomic summaries were created using Quantitative Insights Into Microbial Ecology (QIIME; RRID:SCR_008249). Measures of beta diversity were computed using the Bray Curtis dissimilarity metric (phyloseq, RRID:SCR_013080) in R (R Project for Statistical Computing, RRID:SCR_001905), and tested for whole community differences across groups using permutational multivariate analysis of variance (PERMANOVA) in the adonis command (vegan, RRID:SCR_011950). These results were visualized via Principal Coordinate Analysis (PCoA) ordination (ggplot2, RRID:SCR_014601). Genera which were significantly different between groups (post-adjustment *α*=0.01) was calculated using the Benjamini-Hochberg multiple testing adjustment procedure (DESeq2, RRID:SCR_015687).

### 2.4 Quantification of maternal intestinal short chain fatty acids

SCFA levels were measured in maternal cecal samples by the McMaster Regional Centre of Mass Spectrometry (MR-CMS). Weight equivalent amount of 3.7% HCl, 10μl10 ul of internal standard and 500 μl of diethyl ether was added to each cecal sample, and vortexed for 15 minutes. After vortexing, 400μl of diethyl ether fecal extract was transferred to a clean 1.5 ml Eppendorf tube. In a chromatographic vial containing an insert, 20μl of N-tert-butyldimethylsilyl-N-methyltrifluoroacetamide (MTBSTFA) was added, after which 60μl of diethyl ether fecal extract was added. The mixture was incubated at room temperature for 1 hour and analyzed using GC-MS (Agilent 6890N GC, coupled to Agilent 5973N Mass Selective Detector).

### 2.5 RNA extraction and cDNA synthesis

Maternal intestinal, placental, and fetal small intestinal tissue was homogenized via bead beating in 900μl of TRIzol (Life Technologies, Canada), and centrifuged at 12,000 x g for 10 minutes at 4°C. Supernatant was removed and added to 300μl of chloroform. The solution was thoroughly mixed and incubated at room temperature for3minutes. Samples were centrifuged for 10 minutes at 12,000 x g at 4°C. The aqueous phase of the sample was removed and added to 500μl of isopropanol. Samples were mixed thoroughly, incubated for 20 minutes at room temperature, after which they were centrifuged at 12,000 x g for 10 minutes at 4°C. The supernatant was removed, and the resulting pellet was washed twice with 75% ethanol. RNA was reconstituted in 20μl ultrapure water and quantified using the NanoDrop 2000 spectrophotometer (Thermo Scientific, Canada) with the NanoDrop 2000/2000c software (Thermo Scientific, Canada). The A260/A280 and A260/A230 ratio for all samples were > 2.0 and > 1.5, respectively. All RNA samples were stored at −80°C until experimentation. Due to small amount of starting material, fetal small intestine RNA was extracted using the Qiagen RNeasy Mini Kit as per the manufacturer’s instructions. Following extraction, 2 μg of RNA was used for cDNA synthesis using the Superscript VILO^TM^ cDNA Synthesis Kit (Life Technologies, Canada) as per the manufacturer’s instructions. Each reaction consisted of 4 μl of 5X VILOTM Reaction Mix, 2 μl of 10X SuperScript Enzyme Mix, 4 μl of RNA (concentration of 250ngμl-1) and 12μl of ultrapure water for a total reaction volume of 20 μl. Tubes were incubatedin athermocycler (Bio-Rad C1000 Touch) at 25°C for 10min, followedby42°C for1 hour, and then 85°C for5 minutes. Complementary DNA was stored at-20°C until qPCR assays were performed.

### 2.6 Quantitative PCR

To quantify transcript levels, quantitative PCR was performed using the LightCycler 480 II (Roche) and LightCycler 480 SYBR Green I Master (Roche Diagnostics, 04887352001). Primers were designed using Primer-BLAST software available at NCBI (blast.ncbi.nlm.nih.gov) and manufactured by Life Technologies (Table 1). The following PCR cycling conditions were used for each assay: enzyme activation at 95°C for 5 min, amplification of the gene product through 40 successive cycles of 95°C for 10 sec, 60°C for 10 sec, 72°C for 10 sec. Specificity of primers was tested by dissociation analysis and only primer sets producing a single peak were used. Each plate for a gene of interest contained a standard curve (10-fold serial dilution) generated using pooled cDNA. Each sample and standard curve point was run in triplicate. The crossing point (Cp) of each well was determined via the second derivative maximum method using LightCycler 480 Software, Release 1.5.1.62 (Roche). An arbitrary concentration was assigned to each well based on the standard curve Cp values by the software. The geometric mean of housekeeping genes was determined to use as reference gene value for each intestinal sample. *β*-actin, cyclophilin, and HPRT were used as housekeepers for maternal intestines, cyclophilin and HPRT for fetal intestines, and *β*-actin and UBC for placentas. Relative mRNA levels for each sample were determined by dividing the geometric mean of the triplicate for each sample by that sample’s reference gene value.

**TABLE 1.**
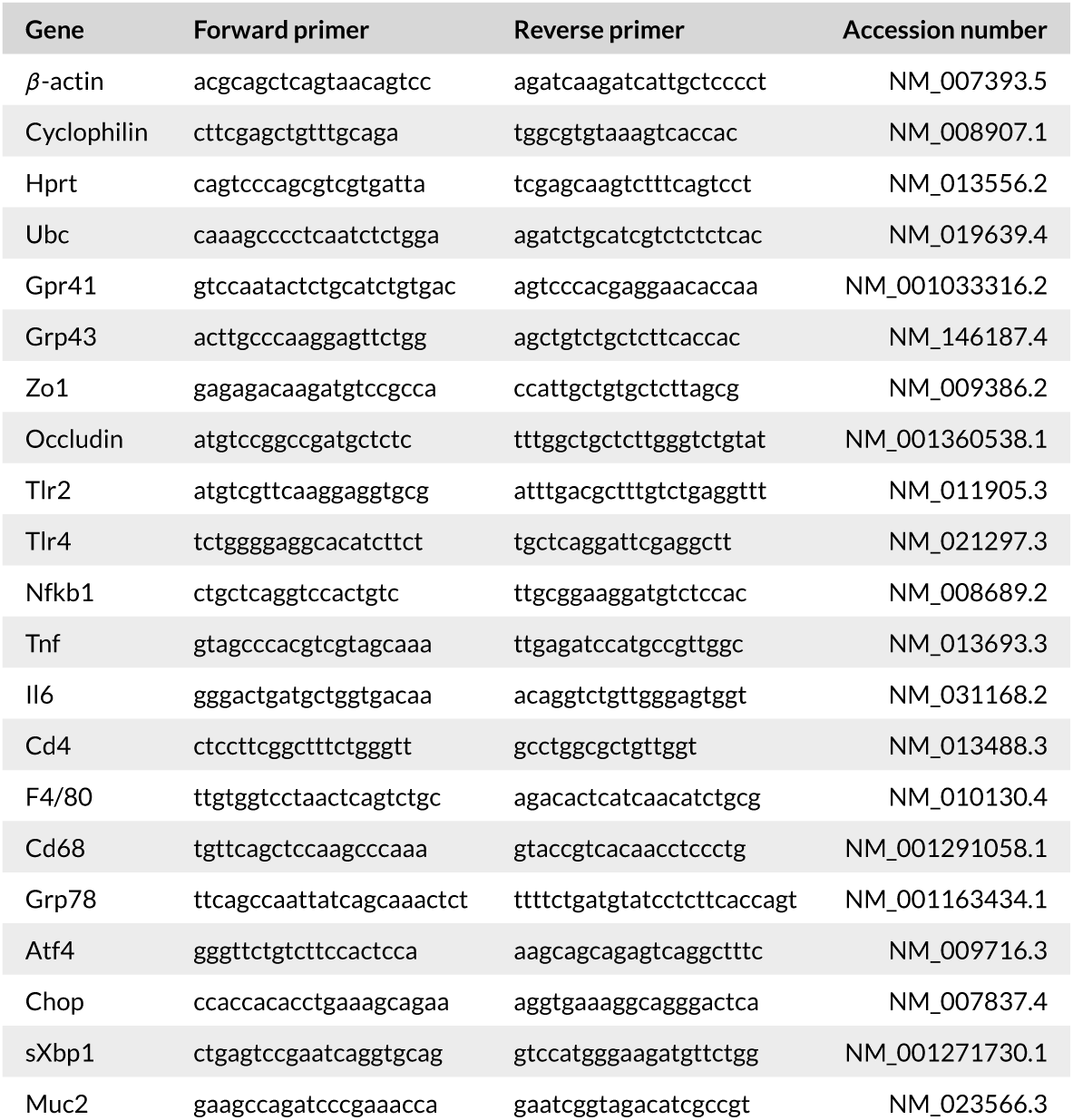
Primer sequences

### 2.7 NF.κB activity and Western Blot analyses

Maternal and fetal intestinal tissue was homogenized with ceramic beads using a Precellys 24 homogenizer at 5 m/s for 60 s in buffer containing 50mMKH2PO4, 5mMEDTA, 0.5mMDTT, and 1.15% KCl with cOmplete protease inhibitor tablets (Roche). Protein concentrations were determined using the bicinchoninic acid (BCA) method in the supernatant after homogenates were centrifuged for 10 min at 10,000 g at 4°C. NF-κB activity was measured in maternal and fetal intestinal homogenates using the TransAM^TM^ NF-κBp65 Transcription Factor Assay Kit according to the manufacturer’s instructions (Cat. no. 40096, Active Motif, Carlsbad, CA, USA).Western blot analyses were performed on ‥ μg of protein using SDS-PAGE. After transfer onto PVDF membrane, blots were blocked for 1 hour at room temperature in 5% BSA TBST and incubated with the following primary antibodies overnight: grp78 (1:2000, Abcam Cat# ab21685, RRID:AB_2119834), phospho-PERK (1:500,Thermo Fisher Scientific Cat# MA5-15033, RRID:AB_10980432), PERK (1:1000, Cell Signaling Technology Cat# 3192, RRID:AB_2095847), phospho-eIF2*α* (1:1000, Cell Signaling Technology Cat# 9721, RRID:AB_330951), eIF2*α* (1:1000, Cell Signaling Technology Cat# 9722, RRID:AB_2230924), occludin (1:1000, Abcam Cat# 168986, RRID: AB_2744671), phospho-JNK (1:500, Abcam Cat# ab124956, RRID:AB_10973183), JNK (1:500, Abcam Cat# ab179461, RRID: AB_2744672), *β*-actin (1:5000, Cell Signaling Technology Cat# 5125, RRID:AB_1903890). Blots were washed in TBST and incubated with HRP-conjugated goat anti-rabbit IgG secondary antibody (Abcam Cat# ab6721, RRID:AB_955447) for 1 hour at room temperature. Blots were developed using BioRad ClarityTM Western enhanced chemiluminescence (ECL; Bio-Rad, 170-5061), images captured using Bio-Rad ChemiDoc MP System and densitometric quantification was carried out using ImageLab Software (Bio-Rad) relative to *β*-actin internal controls.

### 2.8 Maternal intestinal barrier integrity

In a separate cohort of mice, a fluorescein isothiocyanate-dextran (FITC-dextran) assay was used to measure in vivo intestinal permeability to assess maternal intestinal barrier integrity prior to and during pregnancy. In both nonpregnant and pregnant (E18.5) mice, a baseline blood sample was collected via tail vein bleed. Plasma was isolated and stored at 4°C protected from direct light until analysis. Each mouse was administered ‥ μl of 80 mg/ml FITC labeled dextran (molecular mass 4kDa; Sigma Aldrich, St Louis,MO, USA) by oral gavage. After 4 hours, a second post-gavage blood sample was collected via tail vein bleed and plasma was isolated. Fluorescence intensity of plasma was measured on a plate reader (BioTek^®^, Oakville, Canada) with an emission 530 nM and excitation at 485 nM. Intestinal barrier integrity was quantified by subtracting the baseline fluorescence values from the post-gavage fluorescence values in each dam and was expressed as relative fluorescence units (RFU).

### 2.9 Biochemical Assays

#### LPS

An in-house colourimetric reporter bioassay was used for the quantification of NF-κB activation by the pattern recognition receptor TLR-4 in response to LPS (Verschoor et al., 2015). Briefly, an LPS-responsive reporter cell line was generated by transiently transfecting the pNifty2-SEAP plasmid into a commercially available HEK293 cell line (RRID:CVCL_Y406) expressing TLR4, MD2, and CD14 (Invivogen, CA, USA). Cells were seeded at 4 × 10^3^ per well in a 96-well plate in Complete DMEM for 24 hours. Serum samples were diluted 1:5 in phosphate buffer saline (PBS), then 1:1 in sterile water, and heat inactivated at 75°C for 5 min. Complete DMEM media was removed prior to the addition of heat-inactivated plasma in HEK Blue Detection (HBD) Media (Invivogen, CA, USA); 10μl of sample and 190μl of HBD media was added into the appropriate wells. Readings were performed at 630 nm, 72 hours after stimulation, and background levels were subtracted from relative absorbance units. Standard curves generated using different doses of purified LPS were used to calculate the relative amount of LPS in serum.

#### TNF and IL-6

Maternal serum TNF and IL-6 levels were measured using Milliplex Map Mouse Cytokine Magnetic Bead Panel (EMB Millipore, MCYTOMAG-70K, Darmstadt, Germany), that uses the Luminex xMAP detection method. The protocol was followed as per the manufacturer’s instructions.

### 2.10 Immunohistochemistry

Paraformaldehyde (4%) fixed placentas were processed, embedded in paraffin and sectioned at .μm, or at .μm for F4/80 and activated caspase-3 (AC3). To inhibit endogenous peroxidase activity, placental sections were immersed for 10 minutes in 30% aqueous hydrogen peroxide, except for F4/80 and AC3 which were immersed for 25 minutes in 0.5% or 1.5% hydrogen peroxide in methanol respectively. Antigen retrieval was performed by incubating tissues in 10mM sodium citrate buffer with Tween, pH 6.0, at 90°C for 12 min, except for F4/80 where antigen retrieval was performed by incubation with 1 mg/ml trypsin in ddH2O (Sigma-Aldrich T7168) for 12 minutes at 37 degrees Celsius (F4/80). Nonspecific binding was blocked with 5%bovine serum albumin for 10 minutes or 60 minutes (F4/80 and AC3)at room temperature (RT) and sections were incubated overnight at 4°C with the following primary antibodies: CD31(1:200, Abcam Cat# ab24590, RRID:AB_448167), *α*-SMA (1:400, Santa Cruz Biotechnology Cat# sc-32251, RRID:AB_262054), VEGF (1:400, Santa Cruz Biotechnology Cat# sc-152, RRID:AB_2212984), VEGFR2 (1:400, Santa Cruz Biotechnology Cat# sc-6251, RRID:AB_628431), carbonic anhydrase-IX (1:600, Abcam Cat# ab15086, RRID:AB_2066533), F4/80 (1:100, Abcam Cat# ab6640, RRID:AB_1140040), and AC3 (Cell Signaling Technology Cat# 9661, RRID:AB_2341188). The following day, tissue sections were incubated at RT for 2 hours with anti-mouse biotinylated secondary antibody (Sigma-Aldrich Canada Ltd)or for 1 hour with anti-rat (Vector Laboratories BA-4001)or anti-rabbit (Vector Laboratories PK6101) secondary antibody. Biotinylated secondary antibodies were further incubated with ExtrAvidin (Sigma-Aldrich Canada Ltd) or HRP conjugated streptavidin (Vector Laboratories PK6101) for 1 h. Visualization of biotinylated antibody labelling was performed by chromogenic development using 3,3‘-diaminobenzidine (D4293; Sigma-Aldrich), and all sections were counterstained with Gills 2 hematoxylin (GHS216; Sigma-Aldrich). For quantification of F4/80 and AC3 staining, whole tissue sections were scanned using a Nikon Eclipse Ni microscope at 10X magnification and a region of interest was defined around either the labyrinth compartment (F4/80) or total placenta (AC3). Chromogen detection was performed using an automated threshold function using Nikon NIS Image Analysis software (v4.30.02, RRID:SCR_014329). For all other experiments, measurements were taken in six fields of view per section at 250× magnification and image analyses were performed using an Olympus BX-61 microscope and integrated morphometry software (MetaMorph Microscopy Automation and Image Analysis Software, RRID:SCR_002368). All the image analyses were performed by an investigator blind to the study groups.

### 2.11 Nanostring

In a subset of placental samples, a custom nCounter Reporter CodeSet designed by Dr. Ask (McMaster University) was used to analyze placental gene expression using NanoString nCounter gene expression system (Table 3). Briefly, 100 ng of total RNA was incubated overnight at 65°C with nCounter Reporter CodeSet, Capture ProbeSet, and hybridization buffer. Post-hybridization, the samples were processed using the nCounter Prep-station and nSolver Analysis Software 2.5 was used to normalized raw NanoString data to 6 housekeepers: UBC, TUBA1A, HPRT, IPO8, GusB and *β*-actin.

**TABLE 3.**
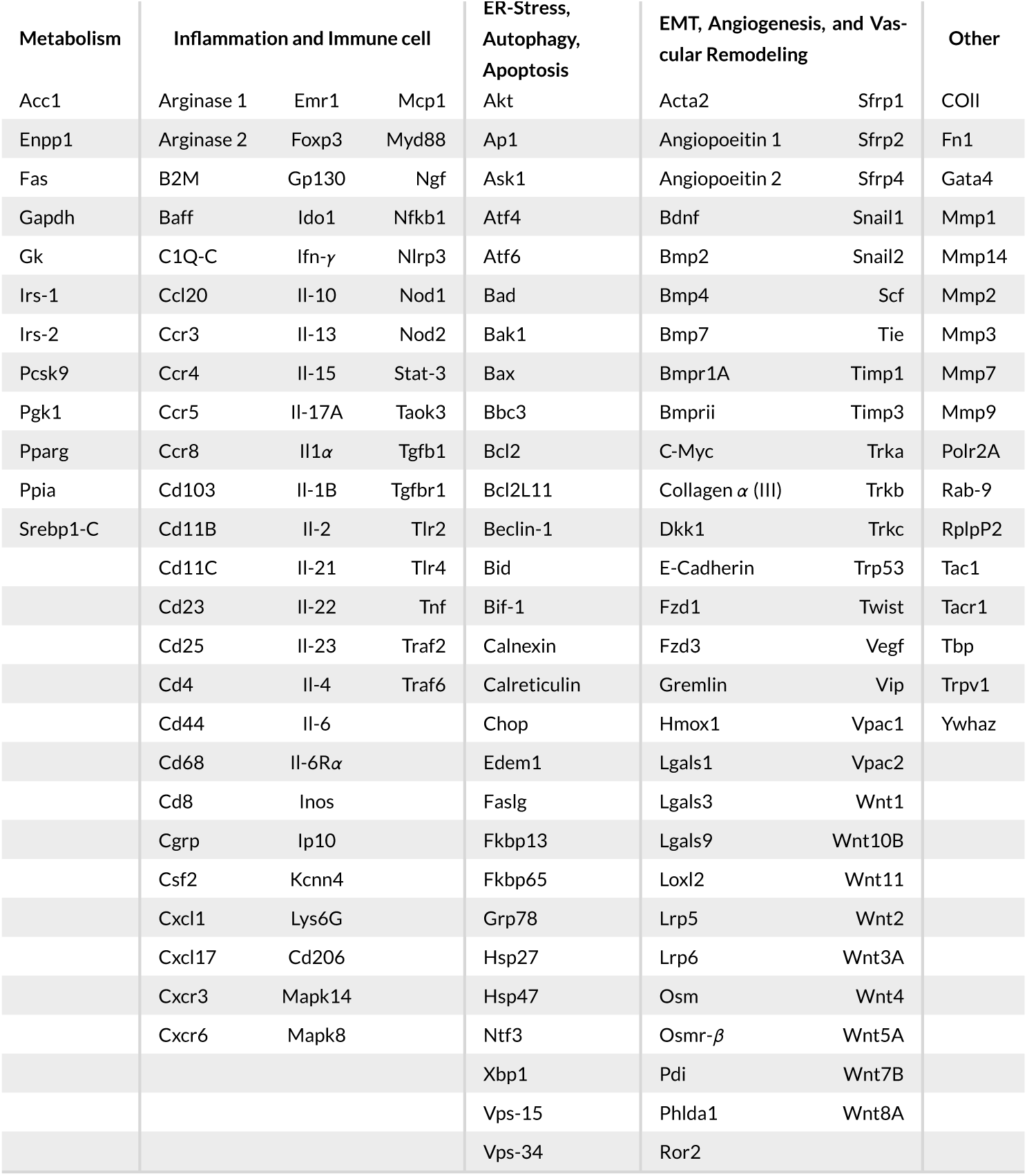
Genes in NanoString nCounter CodeSet excluding housekeepers

### 2.12 Placental metabolomics

Frozen placental tissue samples were lyophilized, powdered and weighed accurately on an electronic balance (3-5 mg). A modified, two-step Bligh Dyer extraction was employed to extract polar, hydrophilic metabolites from the placental tissue. The extraction procedure consisted of adding 64 μl ice cold methanol:chloroform (1:1) followed by 26 μl de-ionized water to enable phase separation. After vortexing for 10 min and centrifugation at 2,000 g at 4°C for 20 min, the upper aqueous layer was obtained. A second extraction on the residual placental tissue was performed by adding another aliquot of 32μl of methanol:deionized water (1:1), followed by the same vortexing and centrifugation procedure described above. The second upper aqueous layer was then transferred and combined with the first aliquot of aqueous layer from the first round of extraction for a total volume of about 80 μl of placental tissue extract. Prior to analysis, 5μl of 3-chloro-L-tyrosine (Cl-Tyr) and 2-naphthalenesulfonic acid (NMS) were added to 20 μl of placental tissue extract resulting in a final concentration of 25 μM for each internal standard when using both positive and negative mode MSI-CE-MS analysis, respectively (Macedo et al., 2017). Untargeted polar metabolite profiling of placental extracts was performed using multisegment injection-capillary electrophoresis-mass spectrometry (MSI-CE-MS) as a high throughput platform for metabolomics based on serial injection of seven or more samples within a single run (Kuehnbaum et al.,2013; Yamamoto et al., 2016). AllMSI-CE-MS experiments were performed on an Agilent G7100A CE system (Agilent Technologies Inc., Mississauga, ON, Canada) equipped with a coaxial sheath liquid (Dual EJS) Jetstream electrospray ion source with heated nitrogen gas to an Agilent 6230 Time-of-Flight Mass Spectrometer (TOF-MS). Separations were performed on a 120 cm long uncoated fused silica capillary (Polymicro Technologies, AZ, USA) with an inner diameter of 50 μm, using an applied voltage of 30 kV at 25°C. The background electrolyte (BGE) consisted of 1 M formic acid with 15% v/v acetonitrile (pH 1.80) in positive ion mode and 50 mM ammonium bicarbonate (pH 8.50) in negative ion mode, for the separation and detection of cationic and anionic metabolites, respectively. Each experimental run consistedof 6sequential injectionsofplacental tissue extracts that were paired based on maternal diet and placental sex. In addition, a quality control (QC) consisting of a pooled placental extract sample (n = 7) was included in each experimental run to assess system drift and overall technical variance (DiBattista et al., 2017). Overall, fifty-one metabolites from placental extracts were consistently detected in majority of specimens analyzed (> 75%) while also having adequate technical precision (average RSD < 40%) based on repeated analysis of the pooled QC throughout the study (Macedo et al., 2017). All experimental data was integrated and analyzed using Agilent MassHunter Qualitative Analysis B.07.00 (RRID:SCR_015040). Integrated ion response ratios were measured for all placental metabolites relative to their corresponding internal standard and subsequently normalized to the dried mass of each placenta analyzed (mg).

### 2.13 Statistical analysis

Maternal gestational weight gain was analyzed using repeated measures two-way ANOVA, with maternal diet and gestational age as factors. Maternal intestinal mRNA levels and NF-κB activity were analyzed by two-way ANOVA with maternal diet and intestinal section as factors. Maternal serum TNF and circulating LPS levels were analyzed using unpaired t-tests with Welch’s correction. Intestinal permeability was analyzed by Mann-Whitney U-test. SCFA concentrations were analyzed by t-tests with Holm-Sidak correction for multiple comparisons. Data used for all t-tests passed the Kolmogorov-Smirnov test for normality. For all analyses of placentas and fetal intestines, the sample size represents the number of litters analyzed where a representative male and female were analyzed from each litter. Placental NanoString data, placental and fetal gut qPCR data, fetal small intestinal NF-κB activity, and placental immunohistochemistry data were analyzed using two-way ANOVA, with maternal diet and fetal sex as factors. Bonferroni post-hoc analyses were used for multiple comparisons where appropriate. For placental metabolomics, supervised multivariate statistical analysis basedonPartial Least Square-Discriminant Analysis (PLS-DA) was applied for variable selection and ranking significant metabolites (variable importance in projection or VIP score > 1.0) associated with placental metabolomic changes as a function of diet and sex (Chong et al., 2018). All data were also analyzed using GraphPad Prism (GraphPad Prism 6.01 for Windows, GraphPad Software, La Jolla California USA, www.graphpad.com).

## 3 RESULTS

### 3.1 Maternal and fetal phenotype

High-fat fed females were significantly heavier than controls throughout gestation (main effect of diet, p=0.011): E0.5 mean diff.=2.59 (95% CI −0.12-5.31); E10.5 mean diff.=3.35 (95% CI 0.63-6.06); E15.5 mean diff.=2.66 (95% CI −0.058-5.38); E18.5 mean diff.=2.53 (95% CI −0.19-5.24; Control n=9, High-fat n=10). At E18.5, maternal fasting blood glucose and serum insulin did not differ between groups (p=0.61 and p=0.54 respectively). Maternal HFD did not detectably alter litter size (p=0.44), fetal weight (p=0.73), or fetal sex ratio (ANOVA, p=0.94; Fig. 1).

**FIGURE 1.**
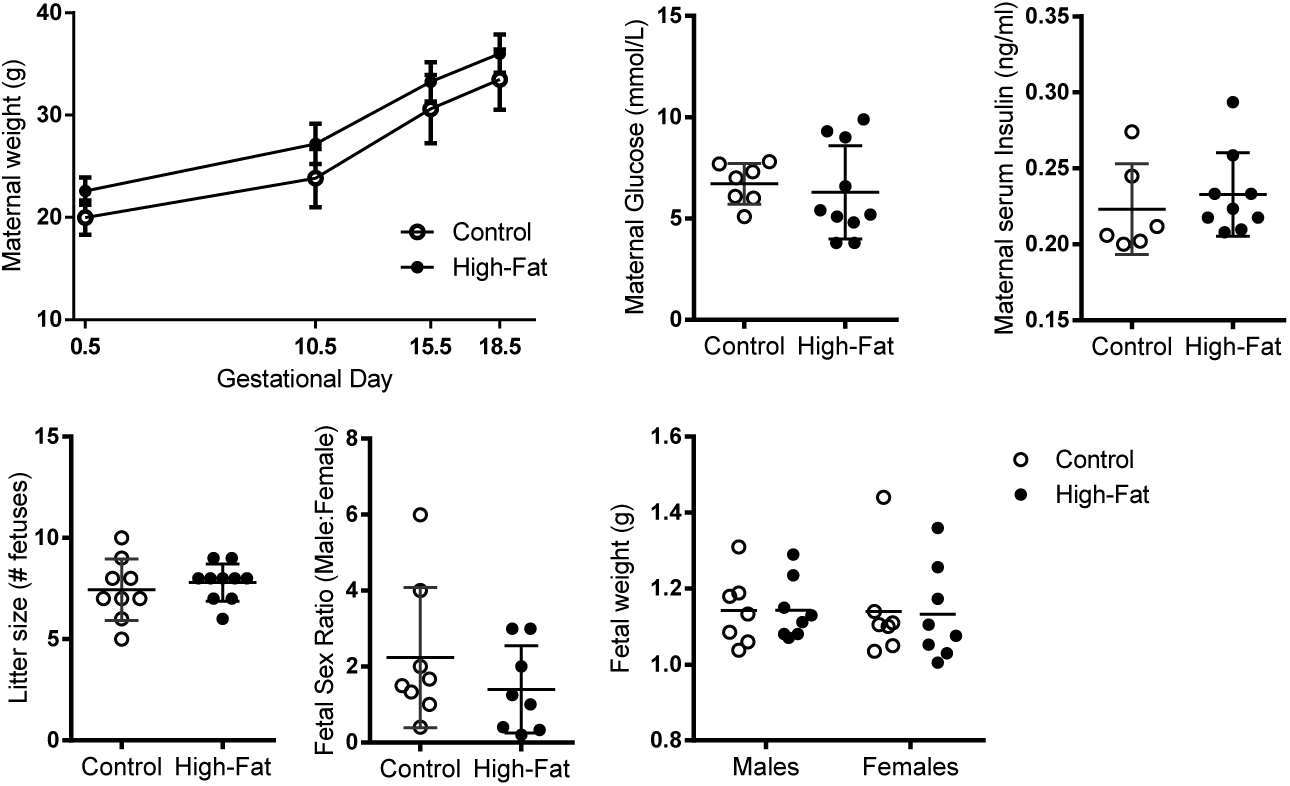
Maternal and pregnancy phenotype. High-fat dams were heavier than controls throughout pregnancy (main effect of diet, p=0.011; main effect of subject (matching), p<0.0001, main effect of gestational day, p<0.0001). Data and mean ± standard deviation. Control, n=6-9; High-fat n=8-10. * p<0.05

### 3.2 Pregnancy and HFD intake induce shifts in intestinal microbiota

At E18.5, the abundance of 29 microbial genera differed significantly between control and high-fat dams (Fig. 2A; Table 2). Community composition of the maternal gut microbiota visualized by Principal Coordinate Analysis (PCoA) using the Bray-Curtis dissimilarity metric shows a distinct separation between control and high-fat samples (Fig. 2B). As we have previously reported (Gohir et al., 2015), the gut microbiota of high-fat fed females was significantly different from control females before pregnancy (adonis: R^2^=0.496, p=0.002) and at E0.5 (R^2^=0.517, p=0.005), E10.5 (R^2^=0.562, p=0.004), E15.5 (R^2^=0.550, p=0.007), and E18.5 (R^2^=0.532, p=0.003).

**TABLE 2.**
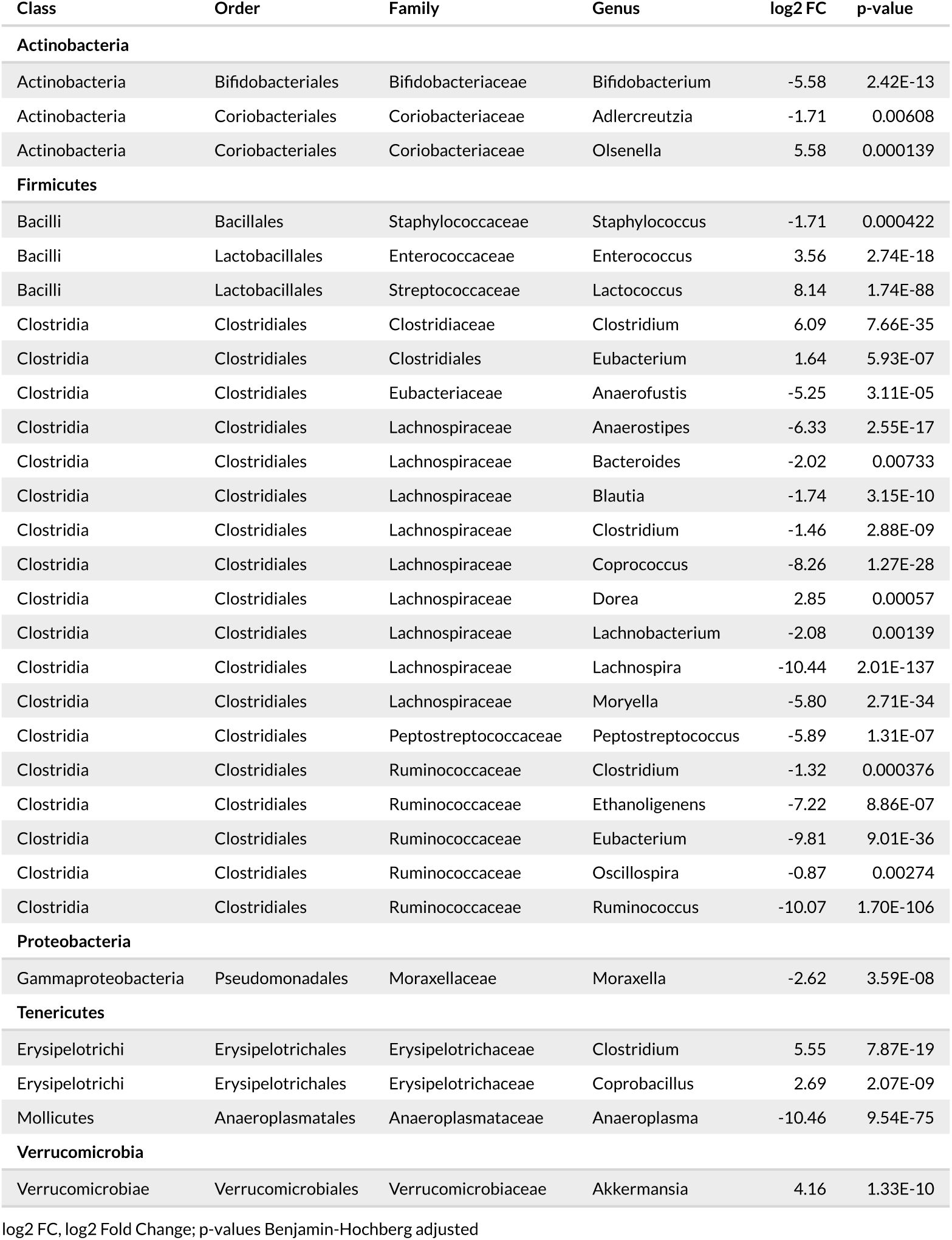
Microbial taxa differing in abundance between control and high-fat damsat E18.5 by DESeq2

| Class | Order | Family | Genus | log2 FC | p-value |
| --- | --- | --- | --- | --- | --- |
| **Actinobacteria** |  |  |  |  |  |
| Actinobacteria | Bifidobacteriales | Bifidobacteriaceae | Bifidobacterium | -5.58 | 2.42E-13 |
| Actinobacteria | Coriobacteriales | Coriobacteriaceae | Adlercreutzia | -1.71 | 0.00608 |
| Actinobacteria | Coriobacteriales | Coriobacteriaceae | Olsenella | 5.58 | 0.000139 |
| **Firmicutes** |  |  |  |  |  |
| Bacilli | Bacillales | Staphylococcaceae | Staphylococcus | -1.71 | 0.000422 |
| Bacilli | Lactobacillales | Enterococcaceae | Enterococcus | 3.56 | 2.74E-18 |
| Bacilli | Lactobacillales | Streptococcaceae | Lactococcus | 8.14 | 1.74E-88 |
| Clostridia | Clostridiales | Clostridiaceae | Clostridium | 6.09 | 7.66E-35 |
| Clostridia | Clostridiales | Clostridiales | Eubacterium | 1.64 | 5.93E-07 |
| Clostridia | Clostridiales | Eubacteriaceae | Anaerofustis | -5.25 | 3.11E-05 |
| Clostridia | Clostridiales | Lachnospiraceae | Anaerostipes | -6.33 | 2.55E-17 |
| Clostridia | Clostridiales | Lachnospiraceae | Bacteroides | -2.02 | 0.00733 |
| Clostridia | Clostridiales | Lachnospiraceae | Blautia | -1.74 | 3.15E-10 |
| Clostridia | Clostridiales | Lachnospiraceae | Clostridium | -1.46 | 2.88E-09 |
| Clostridia | Clostridiales | Lachnospiraceae | Coprococcus | -8.26 | 1.27E-28 |
| Clostridia | Clostridiales | Lachnospiraceae | Dorea | 2.85 | 0.00057 |
| Clostridia | Clostridiales | Lachnospiraceae | Lachnobacterium | -2.08 | 0.00139 |
| Clostridia | Clostridiales | Lachnospiraceae | Lachnospira | -10.44 | 2.01E-137 |
| Clostridia | Clostridiales | Lachnospiraceae | Moryella | -5.80 | 2.71E-34 |
| Clostridia | Clostridiales | Peptostreptococcaceae | Peptostreptococcus | -5.89 | 1.31E-07 |
| Clostridia | Clostridiales | Ruminococcaceae | Clostridium | -1.32 | 0.000376 |
| Clostridia | Clostridiales | Ruminococcaceae | Ethanoligenens | -7.22 | 8.86E-07 |
| Clostridia | Clostridiales | Ruminococcaceae | Eubacterium | -9.81 | 9.01E-36 |
| Clostridia | Clostridiales | Ruminococcaceae | Oscillospira | -0.87 | 0.00274 |
| Clostridia | Clostridiales | Ruminococcaceae | Ruminococcus | -10.07 | 1.70E-106 |
| **Proteobacteria** |  |  |  |  |  |
| Gammaproteobacteria | Pseudomonadales | Moraxellaceae | Moraxella | -2.62 | 3.59E-08 |
| **Tenericutes** |  |  |  |  |  |
| Erysipelotrichi | Erysipelotrichales | Erysipelotrichaceae | Clostridium | 5.55 | 7.87E-19 |
| Erysipelotrichi | Erysipelotrichales | Erysipelotrichaceae | Coprobacillus | 2.69 | 2.07E-09 |
| Mollicutes | Anaeroplasmatales | Anaeroplasmataceae | Anaeroplasma | -10.46 | 9.54E-75 |
| **Verrucomicrobia** |  |  |  |  |  |
| Verrucomicrobiae | Verrucomicrobiales | Verrucomicrobiaceae | Akkermansia | 4.16 | 1.33E-10 |

log2 FC, log2 Fold Change; p-values Benjamin-Hochberg adjusted

**FIGURE 2.**
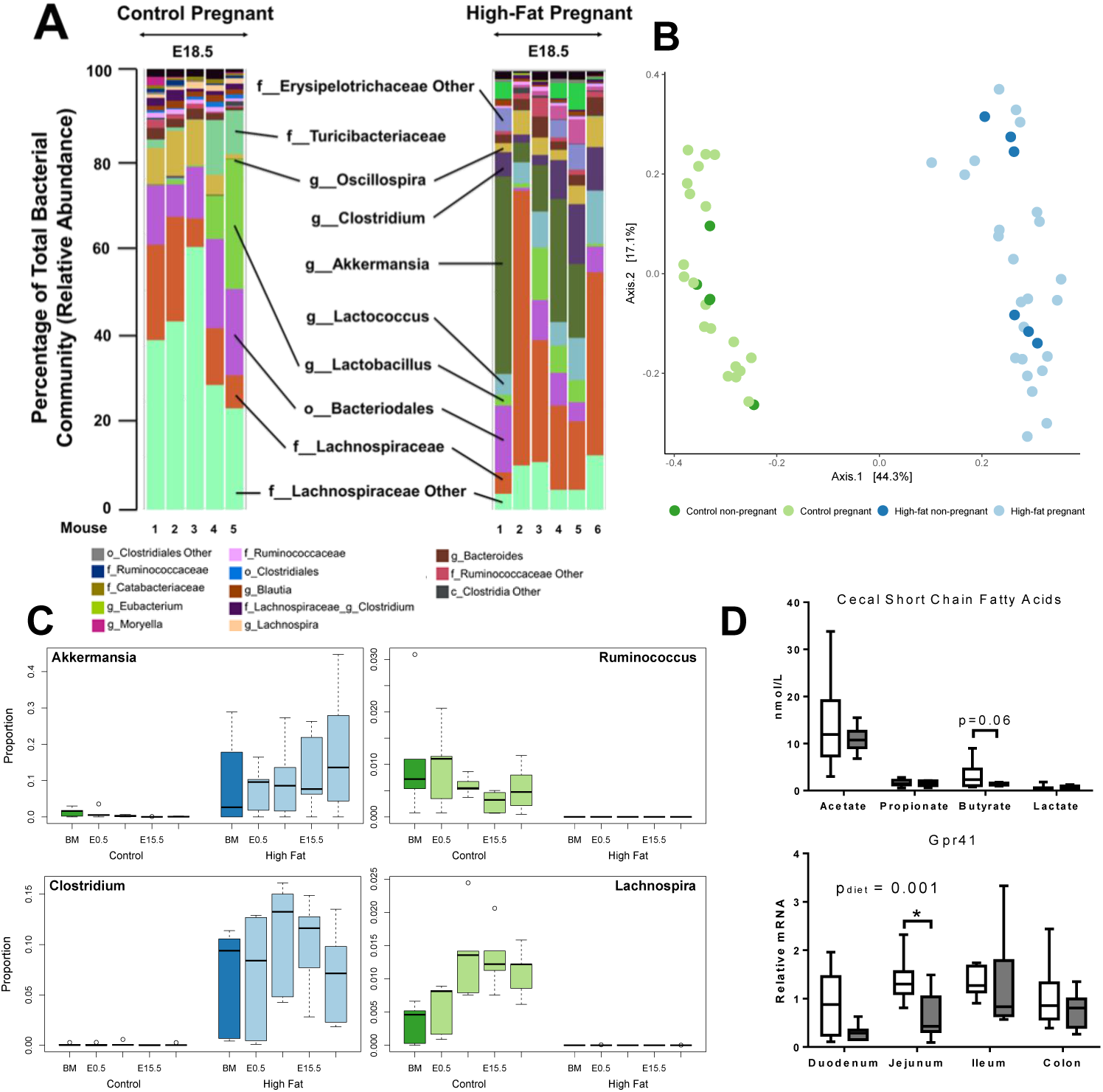
Maternal diet and pregnancy interact to promote shifts in the maternal gut microbiota. A. Relative abundance of the top 25 most abundant taxa for each control (n=5) and high-fat (n=6) dam at E18.5 with significantly different taxa highlighted (f: family, g: genus). B. Principle Coordinate Analysis (PCoA) using the Bray Curtis dissimilarity metric showed distinct separation of gut microbial communities due to maternal diet (adonis p=0.001; C n=5; HF n=6). C. Maternal HFD was associated with increased *Akkermansia* (p=0.0185) and *Clostridium* (p=0.0014), and decreased *Ruminococcus* (p=0.0018) and *Lachnospira* (p<0.0001) relative abundances across all pregnancy time points (Two-way ANOVA;C n=5,HF n=6). D.Cecal SCFAs were measured by GC-MS and relative mRNA levels of SCFA receptors *Gpr41* are shown in in control (open bars, n=9) and high-fat dams (gray bars, n=10). Significance assessed by Bonferroni-adjusted post hoc (* p < 0.05 compared to control).

Consistent with our previous study (Gohir et al., 2015), analysis of whole community differences across all samples found a significant effect of maternal diet (R^2^=0.447, p=0.001) and a small but significant effect of pregnancy timepoint (adonis: R^2^=0.0736, p=0.025). High-fat dams had a higher relative abundance of *Akkermansia*, a mucin degrader, and *Clostridium* (Family *Clostridiaceae*; Fig 2C). The majority of genera within the order *Clostridiales* differing by HFD were decreased (Table 2), including butyrate-producing genera *Lachnospira* and *Ruminococcus* (Morrison and Preston, 2016)(Fig. 2C). Consistent with a decrease in *Lachnospira* and *Ruminococcus*, maternal cecal butyrate levels were also decreased (albeit modestly) in high-fat dams compared to controls at E18.5 (p=0.061; Fig. 2D), as were intestinal transcript levels of SCFA receptor *Gpr41* (main effect of diet p=0.001; Fig. 2D) but not *Gpr43* (data not shown).

### 3.3 HFD alters intestinal inflammation and immune cell marker transcript levels

HFD-induced shifts in the gut microbiota have previously been associated with intestinal macrophage infiltration and inflammation in male mice (Kim et al., 2012). Consistent with this, we found high-fat fed dams had increased intestinal transcript levels of T-cell marker *Cd4* (main effect of diet, p=0.001, Fig. 3A), predominantly in the duodenum and ileum (p=0.036 and p=0.035 respectively). High-fat fed dams had increased transcript levels of monocyte lineage marker *Cd68* (main effect of diet, p=0.0008), predominantly in the duodenum and jejunum (p=0.015 and p=0.024 respectively), while levels of the macrophage maturity marker F4/80 was modestly increased in the ileum (p=0.05; Fig. 3A). To determine whether increases in immune cell markers were associated with intestinal inflammation, we measured transcripts of key pattern recognition receptors involved in immune-related inflammatory signaling. High-fat fed dams had increased intestinal mRNA levels of *Tlr2* (main effect of diet p=0.006) and *Il6* (main effect of diet p=0.035), particularly in the duodenum (p=0.0004 and p=0.018 respectively). *Tlr4* levels were modestly increased in the colon (p=0.05; Fig. 3B). Intestinal NF-κB activation did not differ between high-fat and control dams (main effect of diet p=0.17; data not shown).

**FIGURE 3.**
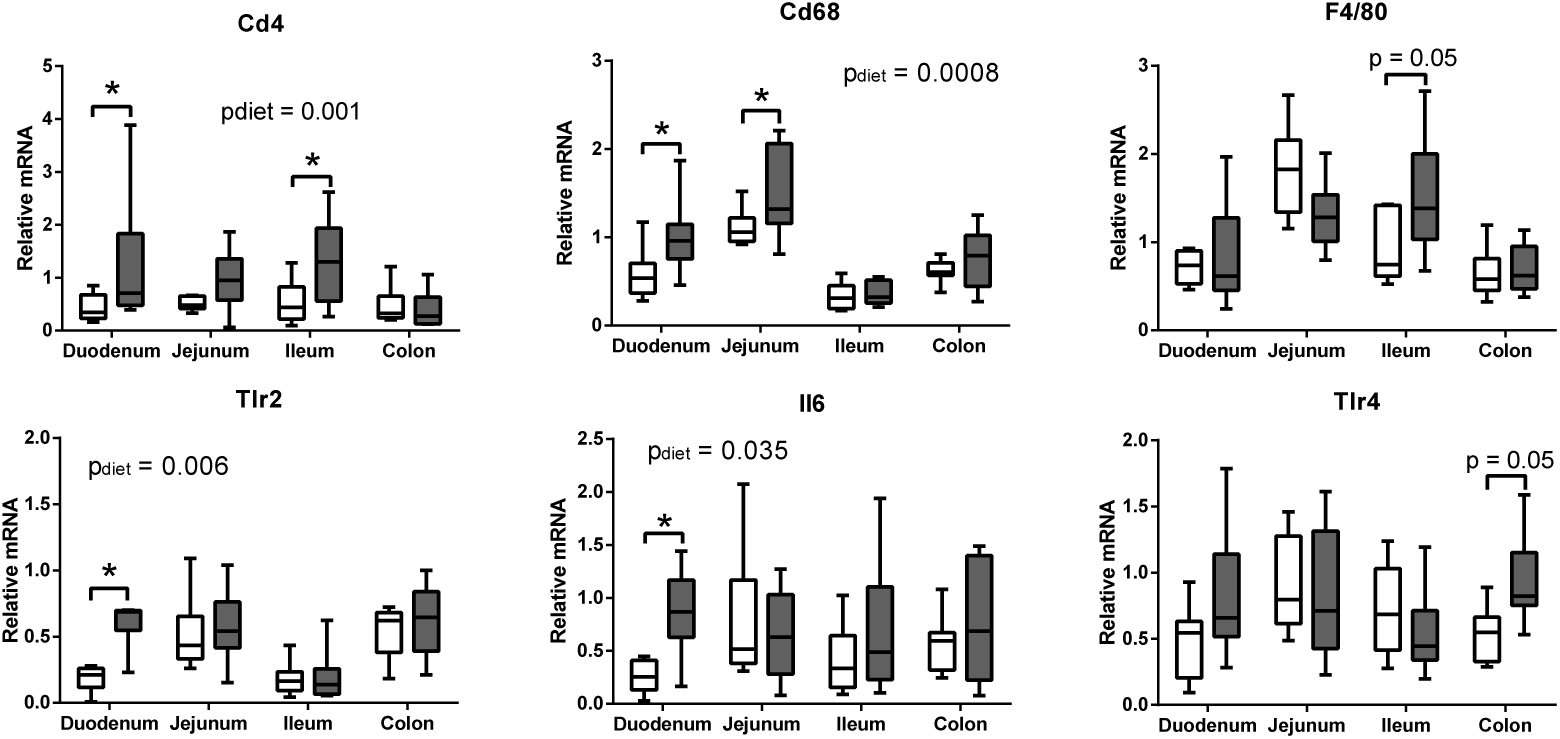
Maternal HFDisassociated with elevated intestinal mRNA levelsofimmune cell markers and inflammatory signaling factors. Relative mRNA levels of A. Cluster of differentiation 4 (*Cd4*), *Cd68*, and F4/80 immune cell markers. B. Toll like receptor (*Tlr*) *2* and *Tlr4*, and interleukin (*Il*) *6*. Data are presented as box and whiskers plots, min to max, where the center line represents the median. Control (open bars, n=9) and high-fat fed dams (gray bars, n=10). Significance assessed by 2-way ANOVA (main effect indicated in text) with Bonferroni post hoc depicted as * p < 0.05 relative to control.

### 3.4 HFD impairs maternal intestinal barrier integrity

To determine whether microbial shifts were associated with functional intestinal changes, we assessed intestinal barrier integrity. Pregnancy increased maternal gut permeability to FITC-dextran (p=0.001) compared to non-pregnant females, and this effect was modestly enhanced in high-fat fed dams compared to controls (p=0.055; Fig. 4). HFD was also associated with decreased transcript levels of tight junction protein zonula occludens-1 (*Zo1*; main effect of diet p=0.0003), most significantly in the ileum (p=0.0027). Occludin mRNA levels however, were increased in the duodenum and ileum (p=0.035 and p=0.0001 respectively, Fig. 4) in high-fat fed dams compared to controls (main effect of diet x section interaction, p<0.0001). Consistent with impaired gut barrier integrity, serum levels of LPS and TNF were higher in high-fat dams compared to controls at E18.5 (p=0.0065 and p=0.024 respectively; Fig. 4).

**FIGURE 4.**
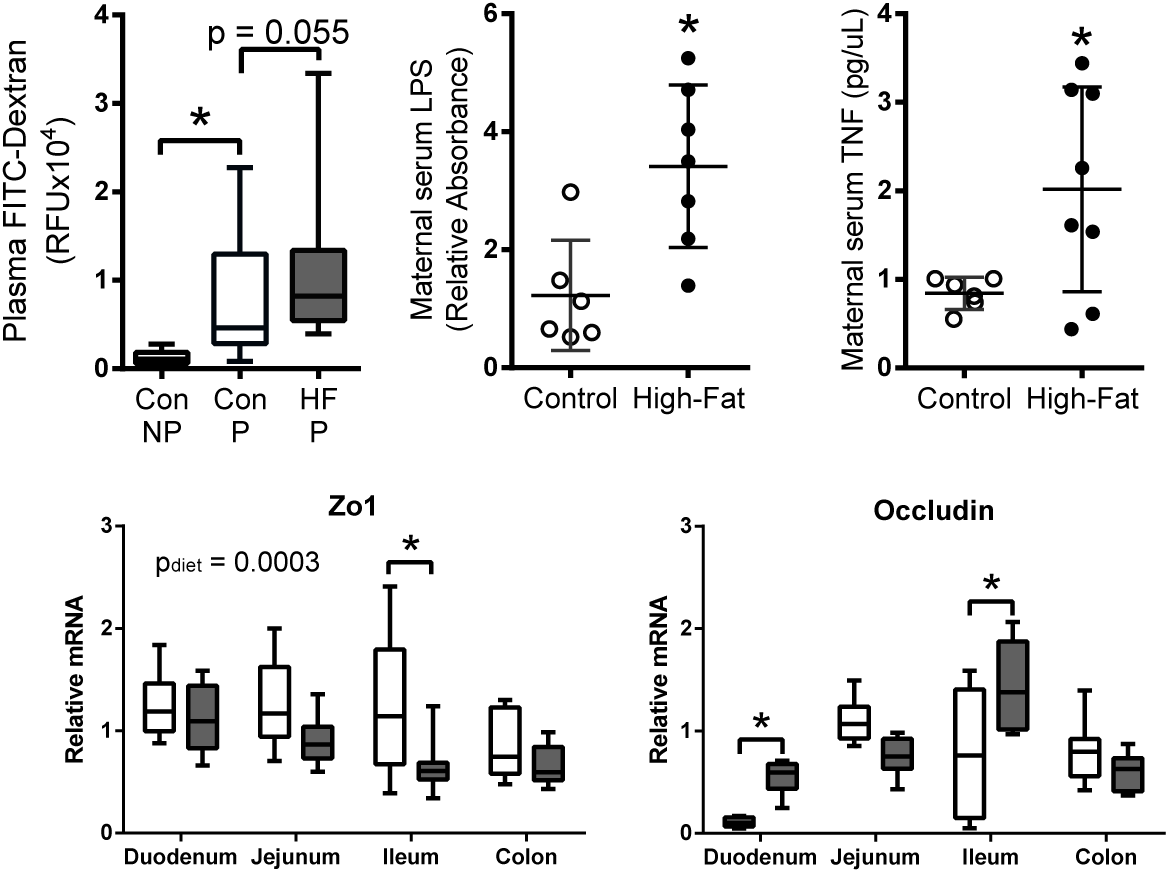
Maternal HFD impairs intestinal barrier integrity and elevates circulating inflammatory markers. FITC-dextran levels are shown in control non-pregnant (Con NP, n=6), pregnant control (Con P,n=14) and pregnant high-fat (HFP, n=14) mice. Maternal serum LPS and TNF levels are shown in control (n=5-6) high-fat dams (n=7-8). Relative mRNA levels of tight junction proteins *Zo1* and Occludin are shown in control (n=9) and high-fat dams (n=10) Data are presented as box and whiskers plots, min to max, where the centre line represents the median. Significance of Bonferroni-adjusted multiple comparisons and t-test analyses depicted as * p < 0.05.

### 3.5 Maternal HFD induces placental hypoxia and apoptosis

Increases in circulating inflammatory factors due to maternal HFD intake result in placental hypoxia (Fernandez-Twinn et al., 2017), which impacts placental vascularization (Hayes et al., 2012; Li et al., 2013), potentially by altering macrophage activation (Zhao et al., 2018). Consistent with these observations, we found increased immunostaining of carbonic anhydrase IX, a placental hypoxia marker, in male (p < 0.0001) and female (p < 0.0001) placentas derived from high-fat fed dams (Fig. 5). This increase was associated with increased immunostaining of vascular endothelial growth factor (VEGF) and its receptor VEGFR2 (Fig. 5). Using immunostaining of an endothelial cell marker (CD31) as a measure of vessel density, we found increased CD31 in high-fat placentas (p<0.0001) and a relative decrease in pericyte-marker alpha-smooth muscle actin (*α*-SMA). A decrease in the CD31: *α*-SMA ratio was used as an indicator of decreased blood vessel maturity (p < 0.0001, Fig. 5). Although placental labyrinth immunostaining of macrophage maturity marker F4/80 was not significantly increased by maternal HFD (main effect of diet, p=0.089), we found increased placental immunostaining of apoptosis-marker activated caspase-3 (AC3) with maternal HFD (main effect of diet, p=0.010; Fig. 5).

**FIGURE 5.**
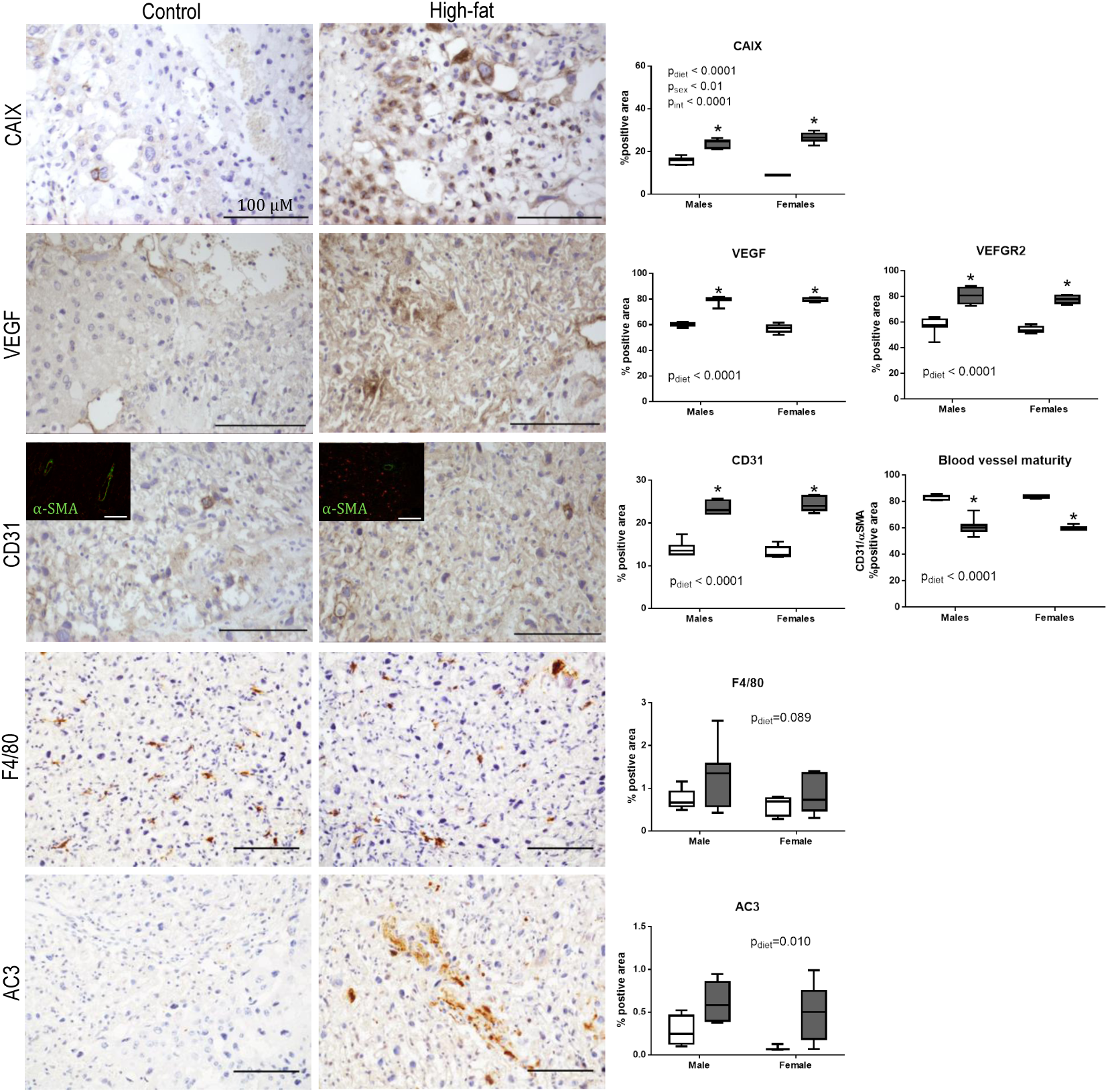
Maternal HFD is associated with placental hypoxia, altered angiogenesis, and apoptosis. Immunostaining of female placental sections from control and high-fat dams. Males show similar staining patterns (photos not shown). Scale bars represent 100 μm. Image analyses of immunostaining show increased carbonic anhydrase (CAIX), VEGF, VEGFR2, and CD31 in placentas from high-fat dams (gray bars, n=5) compared to controls (open bars, n=5). Blood vessel maturity was assessed via the ratio of CD31 (endothelial cell marker) to smooth muscle actin (*α*-SMA, pericyte marker) in green (inset fluorescence staining image). Results of computerized image analyses of immunostaining of F4/80 and activated caspase-3 (AC3) in placentas from control (open bars, n=3-5) and high-fat dams (gray bars, n=5-7). Data are presented as box and whiskers plots, min to max, where the center line represents the median. Significance assessed by Bonferroni-adjusted 2-way ANOVA where *p<0.05. All immunohistochemical assays were performed with negative controls (absence of primary antibodies) and showed no staining (data not shown).

### 3.6 Maternal HFD alters markers of placental growth and development

To explore modulators of placental hypoxia in an untargeted manner, we quantified transcript levels of 178 genes involved in ER stress, autophagy and inflammation pathways in a subset of placentas using a NanoString nCounter system (Table 3).

Shifts in placental macrophages, neutrophils and T cells have previously been associated with pregnancy complica-tions, including preeclampsia (Wallace et al., 2014). Both maternal diet (p=0.018) and sex (p=0.02) were associated with increased transcript levels of *Cd4*, particularly in female placentas (Fig. 6A), and a modest increase in *Cd25* mRNA levels (diet p= 0.07). Increases in these T cell markers were consistent with a significant effect of diet (p =0.017) and modest effect of sex (p=0.06) on interleukin-10 (*Il10*) and modest increases in *Il22* mRNA levels (Fig. 6A). The F4/80 macrophage maturation marker transcript was elevated the placentas of high-fat dams (diet p=0.010, sex p=0.016, Fig. 6A). Reports show that Cd4+ Cd25+ T cells may induce M2 macrophages through arginase and IL10 pathways (Liu et al., 2011), however, Arginase 1mRNA levels were suppressed by maternal HFD and although *Nos2* mRNA levels were unchanged (data not shown), the placental *Arg1*:*Nos2* ratio was significantly lower in high-fat pregnancies (p=0.046; Fig. 6A).

**FIGURE 6.**
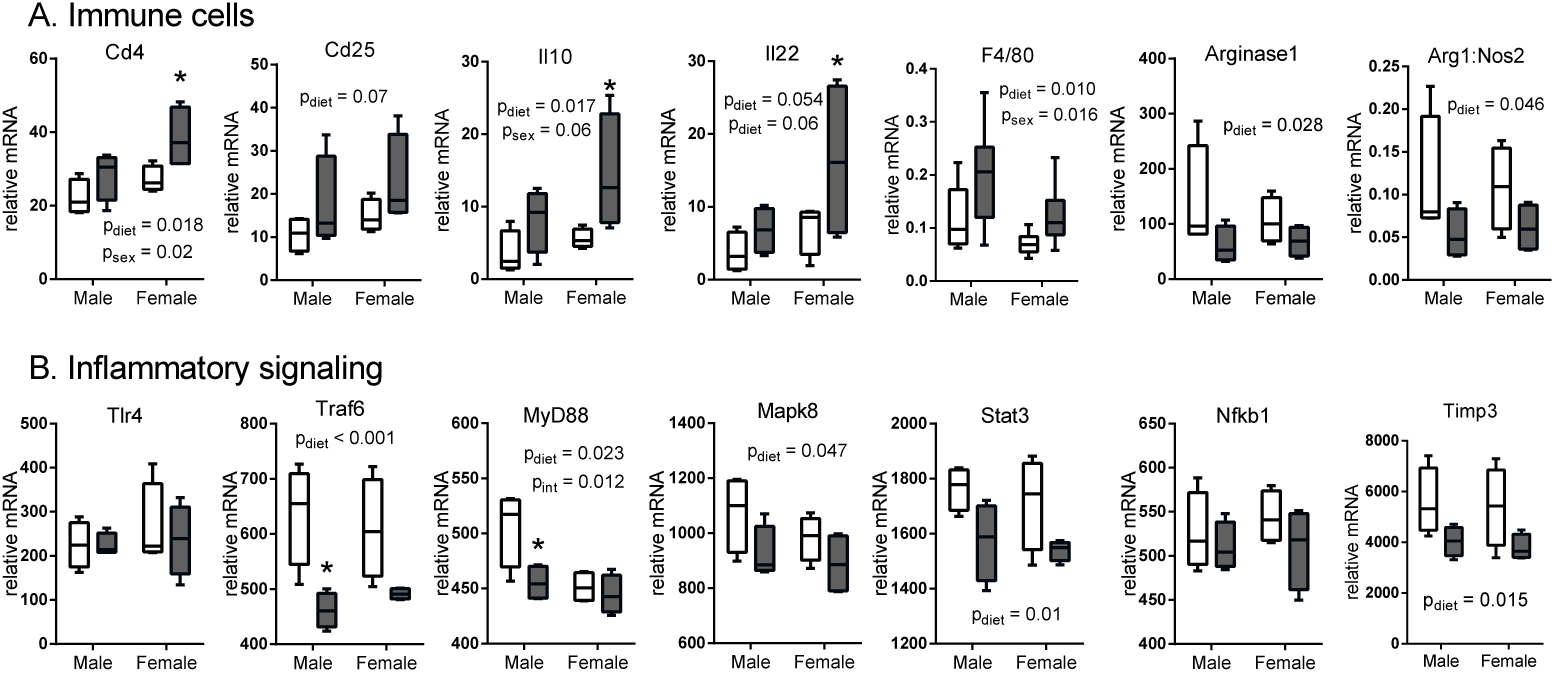
Maternal HFD alters placental mRNA levels of immune cell markers and inflammatory signaling molecules. A subset of placental samples was analyzed using the NanoString nCounter gene expression system for key transcript levels of markersofA.Immune cells and B. Inflammatory signaling factors from control (open bars, n=4) and high-fat dams (gray bars, n=4). Data are presented as box and whiskers plots, mintomax, where the center line represents the median. Significance assessed by Bonferroni-adjusted 2-way ANOVA (main effect indicated as text in graph, post hoc indicated by * p < 0.05 compared to control). CD: cluster of differentiation, F4/80: macrophage maturity marker, IL: interleukin, TLR4: toll like receptor 4, Traf6: TNF receptor-associated factor, MyD88: Myeloid differentiation primary response 88, MAPK8: Mitogen-Activated Protein Kinase 8, STAT3: Signal transducer and activator of transcription 3; NF-κB: nuclear factor kappa-light-chain-enhancer of activated B cells, TIMP3: Tissue Inhibitor of Metalloproteinase 3.

Placental transcript levels of inflammatory receptors were largely unaltered by maternal HFD, including Toll like receptor (*Tlr*) *2* (data not shown), and *Tlr4* (Fig. 6B). Notably however, placental adapter protein transcript levels were significantly decreased by maternal HFD, including tumor necrosis factor receptor (TNFR)-associated factor 6 (*Traf6*), myeloid differentiation primary response 88 (*MyD88*), *Mapk8* and *Stat3* (Fig. 6B), without any changes to pro-inflammatory transcript levels of *Nfkb1* (Fig. 6B).

Neurotrophins (including Neurotrophin 3, *Nt3*) and their tyrosine kinase receptors (specifically *TrkC*) are present at term in cyto-and syncytiotrophoblast cells as well as in the decidua (Casciaro et al., 2009). Immune cells (B and T lymphocytes and monocytes) secrete NT3 and express TrkC, and it has been suggested that placental NT3 modulates inflammatory cell migration to the placenta (Casciaro et al., 2009). We found that maternal HFD intake was associated with a modest increase in *Nt3* transcript levels (diet p=0.06, Fig. 7A) and a significant increase in levels of its receptor *TrkC* (diet p<0.001; Fig. 7A). This is consistent with our observations that *Cd4* and *Cd25* transcripts were increased in high-fat placentas, and increased immunostaining of F4/80 in placental sections.

**FIGURE 7.**
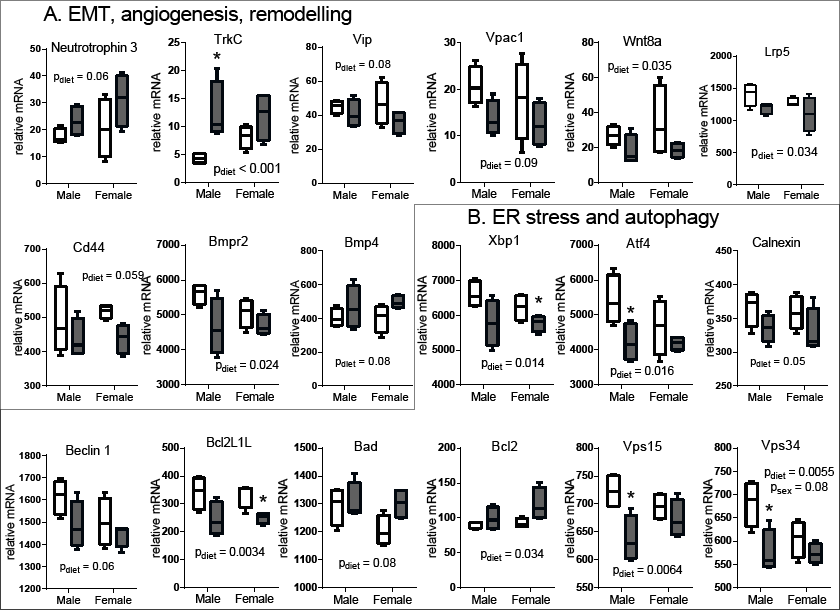
Maternal HFD alters placental mRNA levels of angiogenesis markers and ER stress factors. A subset of placental samples was analyzed using the NanoString nCounter gene expression system for key transcript levels of markers of EMT, angiogenesis, ER stress, UPR and apoptosis from control (open bars, n=4) and high-fat dams (gray bars, n=4). A. Epithelial Mesenchymal Transition (EMT), angiogenesis and tissue remodeling markers. B. Endoplasmic reticulum (ER) stress, Unfolded Protein Response (UPR) and apoptosis and autophagy markers. Data are presented as box and whiskers plots, min to max, where the centre line represents the median. Main effect p values are written as text in the figure. Significance assessed by Bonferroni-adjusted 2-way ANOVA (* p < 0.05). TrkC: Tropomyosin receptor kinase C, CD: cluster of differentiation, VIP: Vasoactive intestinal peptide, VPAC1: Vasoactive intestinal polypeptide receptor 1, BMPR2: Bone morphogenetic protein receptor type II, BMP4: Bone morphogenetic protein 4, LRP5: Low-density lipoprotein receptor-related protein 5, Wnt8a: Wnt family member 8a, XBP1: X-box binding protein 1, ATF4: Activating transcription factor 4, VPS15/34: Serine/threonine-protein kinase, Bcl2L1L: Bcl-2-like 1, Bcl2: B-cell lymphoma 2.

Placental hypoxia is often associated with poor implantation and impairments in epithelial-mesenchymal transition (EMT) and maternal vessel remodeling early in pregnancy (Aplin, 2000; Davies et al., 2016). Transcript levels of factors associated with EMT, cell migration, invasion and vascular remodeling were decreased in placentas from high-fat pregnancies, including *Cd44*, vasoactive intestinal peptide (*Vip*) and its receptor *Vpac1*, bone morphogenetic protein receptor 2 (*Bmpr2*) but not *Bmp4*, as well as WNT signaling molecules *Wnt8a*, and *Lrp5*, (Fig. 7A). These decreases are consistent with our observation of placental hypoxia and reduced *α*SMA immunostaining in high fat placental tissue.

Transcript levels of ER stress factors X-box binding protein 1 (*Xbp1*), and Activating transcription factor 4 (*Atf4*), and the molecular chaperone calnexin, were decreased with maternal HFD intake (Fig. 7B). Consistent with these reductions, mRNA levels of *Vps31* and *Vps34*, which regulate the clearance of protein aggregates, and Beclin-1, a central mediator in autophagy (Kang et al., 2011), were also decreased with maternal HFD (Fig. 7B). Transcript levels of pro-survival *Bcl2L1L* were also decreased and pro-apoptotic *Bad* and *Bcl2* were increased in placenta from high-fat pregnancies (Fig. 7B).These increase sin Bad and Bcl2 are consistent with our observed increases in AC3 immunostaining in the placenta.

### 3.7 Maternal HFD alters placental metabolomics

Recently, the placental metabolome has become a topic of great interest as changes in the relative concentrations of metabolites occurinsome complicated pregnancies (Fattuoni et al., 2018). Weanalyzedfifty-one polar/ionic metabolites in placental extracts normalized to dried tissue weight using multisegment injection-capillary electrophoresis-mass spectrometry (MSI-CE-MS). The metabolomes of high-fat and control placenta were well discriminated by PLS-DA 2D (Fig.8A). We observed slightly more biological variation in metabolite levels in control placentas (median RSD = 57%; females= 78%, males= 35%) compared to high-fat (median RSD = 42%; females: 59%, males = 26%), whereas technical precision was acceptable based on repeated analysis of a pooled placental quality control extract (median RSD = 15%). The Variable Importance in Projection (VIP) scores plot (Fig. 8B) summarizes the top metabolites driving the separation between high-fat and control placentas. Maternal HFD was associated with decreased levels of free carnitine (C0; main effect of diet p=0.0006), short-chain and medium-chain acylcarnitines (C2, C3, C8 and C10; main effect of diet p=0.0007, p=0.0013, p=0.0026, and p=0.0034 respectively), and trimethylamine-N-oxide (TMAO; main effect of diet p=0.0046).

**FIGURE 8.**
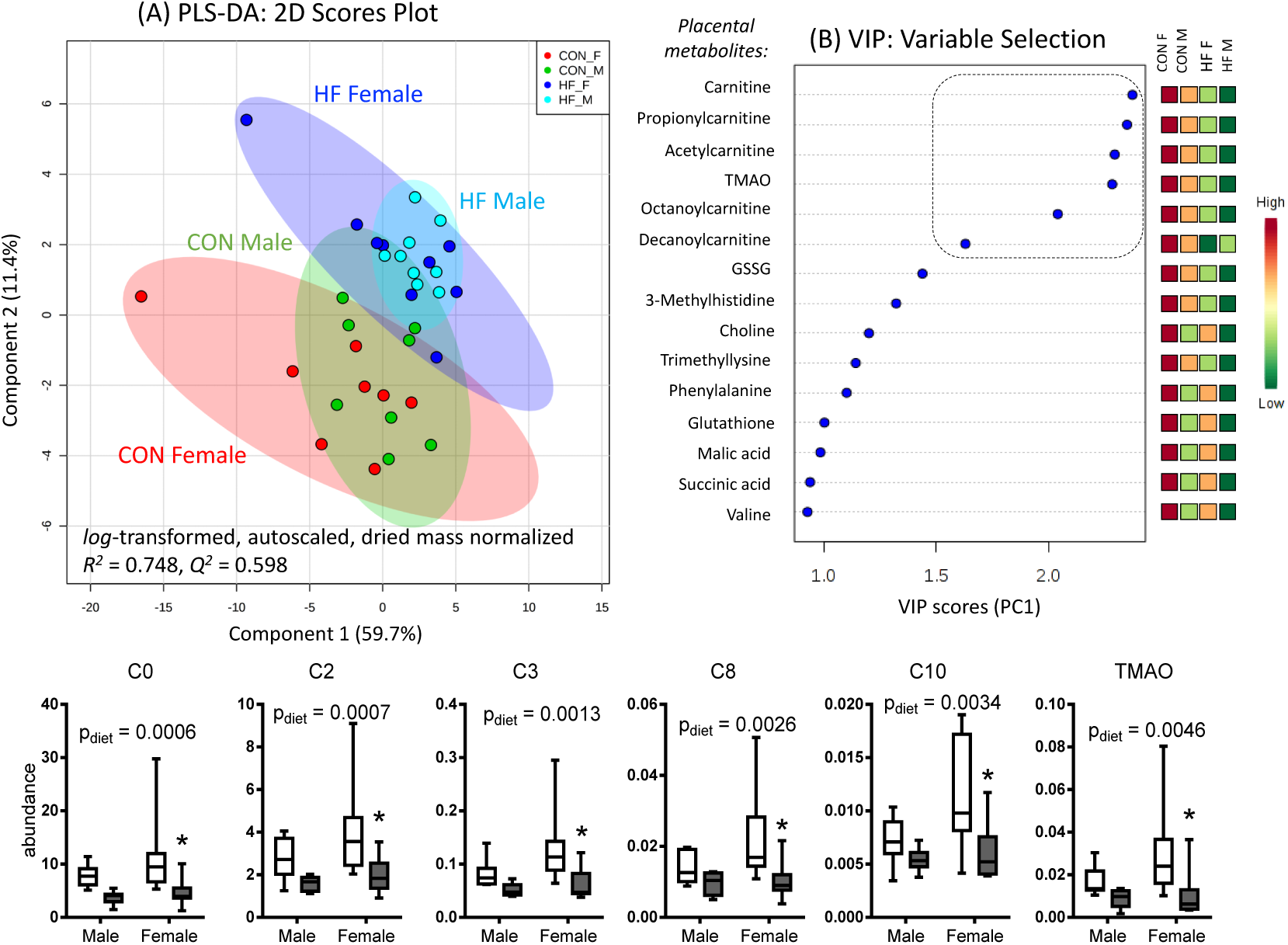
Maternal HFD is associated with decreased placental carnitine, and its related short/medium-chain acylcarnitine derivatives and TMAO. PLS-DA 2D scores plot illustrating diet and sex dependent changes in the placental metabolome of control female (CON_F red, n=8) and control male (CON_M green,n=8), ascomparedtoplacental extracts from high-fat female (HF_F, blue, n=9) and high-fat male (HF_M, light blue, n=9) groups. Supervised multivariate data analysis demonstrated good model accuracy (R^2^ = 0.75) and robustness (Q2 = 0.60) when using leave one-out-at-a-time cross validation. A variable importance in projection (VIP) along PC 1 was used for variable selection, which summarizes the top 15-ranked placental metabolites primarily associated with high-fat maternal diet (VIP > 1.0). The dashed box within VIP plot highlights that 6 placental metabolites were found to be significantly lower in high-fat placentas when using a FDR adjustment (q < 0.05). Data are presented as box and whiskers plots, min to max, where the center line represents the median. Significance assessed by Bonferroni-adjusted 2-way ANOVA and depicted as * p > 0.05. C0: Carnitine; C2: Acetylcarnitine; C3: propionylcarnitine; C8: octanoylcarnitine; C10: decanoylcarnitine; TMAO: Trimethylamine-N-oxide.

### 3.8 Maternal HFD impacts fetal intestinal cell stress pathways

As maternal HFD has previously been associated with increased fetal intestinal inflammation (Yan et al., 2011), we measured NF-κB activation using a TransAm p65 kit, which measures the activated form of the p65 subunit. We found that fetal intestinal NF-κB activation was increased by maternal HFD in the small intestine (main effect of diet p=0.0007), particularly in female fetuses (p=0.0031; Fig.9A). NF-κB activation was also increased in the large intestine of female (p=0.026), but not male (p>0.99) fetuses (Fig. 9A). This was not associated with increased mRNA levels of *Tnf*, *llfβ II10* in either the small or large intestine (data not shown).

**FIGURE 9.**
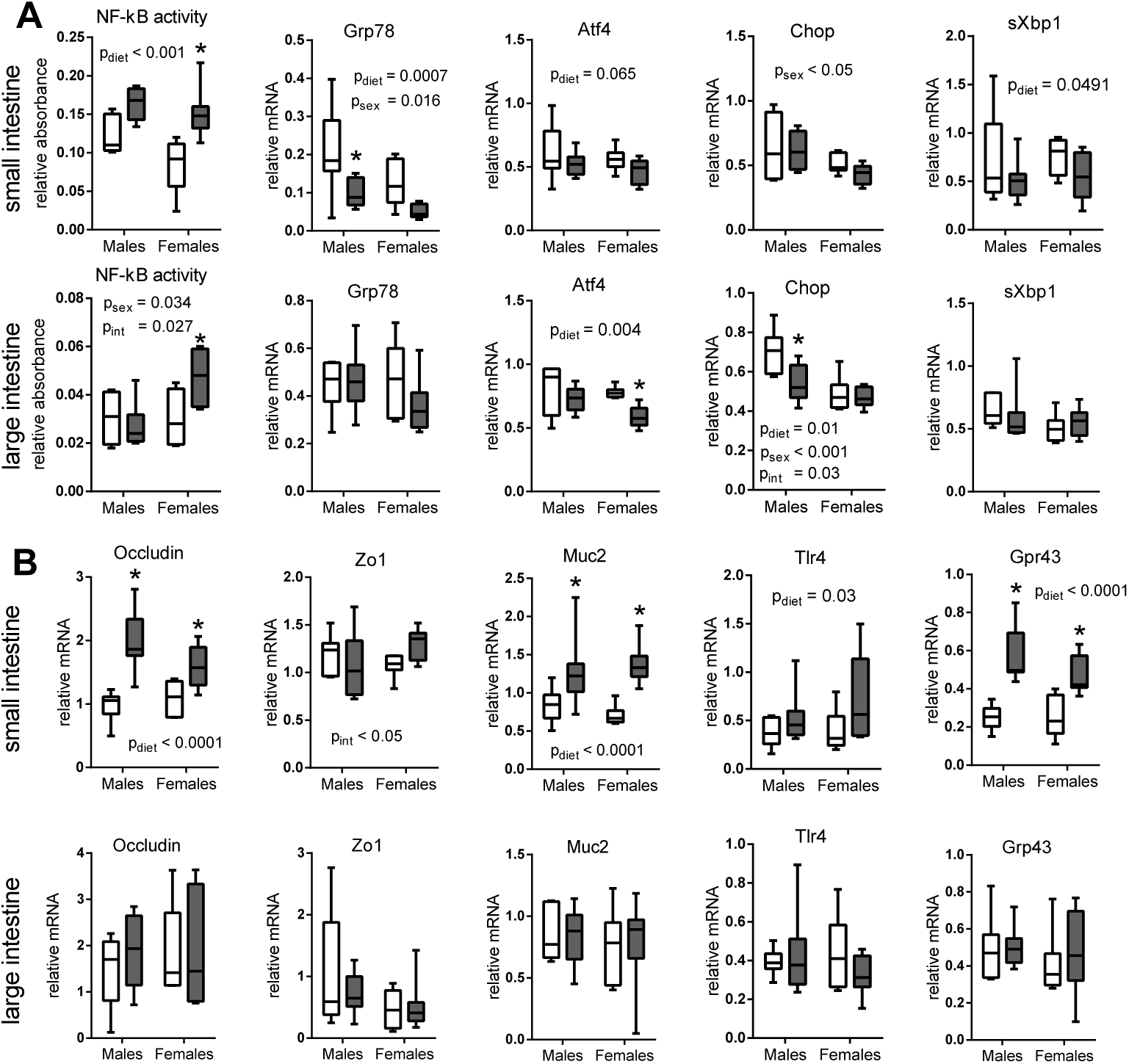
Transcript levels of factors regulating fetal gut barrier, UPR, and NF-κB activity are altered by maternal HFD. A. In fetal small intestines, mRNA levels of *sXBP1* and *Grp78* were decreased in association with maternal HFD. In fetal large intestines, mRNA levels of *Atf4* and *Chop* were decreased in association with maternal HFD. B. In fetal small intestines, *Muc2*, occludin, and *Tlr4* mRNA levels were increased in association with maternal HFD, while there was no significant effect of maternal HFD on mRNA levels of *Zo-1*. NF-κB activation was increased with maternal HFD in fetal small intestines and female fetal large intestines. Data are presented as box and whiskers plots, min to max, where the center line represents the median. Main effect p values are written as text in the figure. Small intestine (n = 7-8) and large intestine (n = 6-8) in control (open bars) and high-fat (gray bars) female and male fetuses [small intestine n = 5-7, large intestine n = 4-7 for NF-κB].Significance assessed by Bonferroni-adjusted 2-way ANOVA and depicted as * p < 0.05.

As NF-κB activation is influenced by ER stress, we explored ER-stress pathways in the fetal intestine. ER-chaperone 78 kDa glucose-regulated protein (*Grp78*) mRNA was decreased in the small intestine (main effect of diet p=0.0007) but not in the large intestine in fetuses exposed to maternal HFD (Fig. 9A), but protein levels were unchanged (data not shown). Maternal HFD was also associated with decreased *sXbp1* mRNA levels in fetal small intestine but not large intestine (main effect of diet p=0.0491 and p=0.98 respectively). Although no difference in PERK or eIF2*α* phosphorylation was detected by western blot (main effect of diet p=0.57 and p=0.46 respectively, data not shown), maternal HFD decreased transcript levels of *Atf4* in fetal small and large intestine (p=0.065 and p=0.004 respectively) and CCAAT/-enhancer-binding protein homologous protein (*Chop*) in fetal large intestine (main effect of diet p=0.01) particularly in male fetuses (p=0.003; Fig. 9B).

To investigate whether maternal HFD was associated with altered fetal intestinal barrier development, we quantified mRNA levels of tight junction components. Maternal HFD increased occludin mRNA levels (main effect of diet p < 0.0001), but not protein levels (main effect of diet p=0.33, data not shown) in the fetal small intestine. There was no detectable effect of maternal diet on *Zo1* mRNA levels in the fetal small or large intestine (main effect of diet p=0.57 and p=0.53 respectively). Maternal HFD also increased transcript levels of mucin 2 (*Muc2*), *Grp43*, and *Tlr4* in the fetal small intestine (p<0.0001, p<0.0001, and p=0.032 respectively; Fig. 9B).

## 4 DISCUSSION

Consistent with our previous work (Gohir et al., 2015), we found that HFD intake resulted in a decreased relative abundance of SCFA producing bacterial genera. We now show this decrease to be accompanied by lower maternal cecal butyrate levels, impaired maternal intestinal barrier integrity, and increased circulating LPS and TNF levels in high-fat fed dams. We show maternal HFD induced placental hypoxia and impaired placental vascularization, and that high fat diet intake was associated with a changes in placental metabolomics strongly influenced by a decrease in carnitine levels. Finally, we extend previous observations that maternal obesity results in fetal gut inflammation (Yan et al., 2011) to include changes in key factors involved in the UPR and increased NF-κB activation in the fetal intestine. The shifts in pregnant microbiota due to HFD and pregnancy found here are similar to our previous results (Gohir et al., 2015) and may contribute to metabolic adaptations to pregnancy (Koren et al., 2012), although his has not yet been explicitly proven.

Short chain fatty acids (SCFAs) produced by anaerobic bacteria play a key role in regulating intestinal immunity (Smith et al., 2013). Despite the widespread interest in SCFAs and GPRs and their role in obesity and inflammation, our understanding of their function remains unclear. DataonGPR41 and 43 knockout models remains inconsistent, showing both inflammation promoting and inflammation resolving actions (Ang and Ding, 2016). Despite these inconsistencies, SCFA supplementation to high fat diet induced obese mice suppressed diet induced weight gain, attenuated HFD induced increases in GPR transcripts in the colon and promoted changes in bacterial taxa, particularly *Lachnospiraceae* (Lu et al., 2016). In our study, high-fat fed dams had a decreased relative abundance of thirteen bacterial genera belonging to the butyrate-producing families *Lachnospiraceae* and *Ruminococcaceae*, and this was accompanied by a modest decrease in cecal butyrate. We further found a decreased relative abundance of *Bifidobacterium*, which is known to promote butyrate production through cross-feeding (Rivière et al., 2016) and has previously been associated with endotoxemia in male diet-induced obese mice (Cani et al., 2008). These microbial shifts were associated with decreased transcript levels of the SCFA receptor *Gpr41* in intestines of high-fat dams.

Shiftsinthe microbiota due to HFD are often accompanied by gut inflammation and macrophage infiltration (Kim et al., 2012). The increases we found with HFD in transcript levels of monocyte lineage marker *Cd68* and macrophage maturity marker F4/80 are consistent with increased macrophage infiltration. Increased mRNA levels of *Cd4* may be attributed to T cells, monocytes, macrophages, or dendritic cells and although it is not possible to determine the specific cell type associated with this increase, this may be reflective of an increase in tissue resident macrophages (Shaw et al., 2018). Increased duodenal mRNA levels of *Tlr2* and *Il6* are consistent with intestinal inflammation in high-fat dams (Hausmann et al., 2002). Future studies will investigate the maternal gut histologically and to localize proteins in order to fully understand impacts on maternal intestinal function. Unfortunately, this was beyond the scope of the present study.

We found that high-fat fed dams had modestly increased gut permeability to FITC-dextran. The increased intestinal permeability we observed with pregnancy in control mice suggests that decreased intestinal barrier integrity may be a normal adaptation to pregnancy, and that maternal HFD may exacerbate this intestinal adaptation. Previous work suggests that increased paracellular permeability may be due to decreased expression of tight junction proteins (Al-Sadi et al., 2011; Lee, 2015). Consistent with this, maternal HFD intake was associated with decreased *Zo1* transcript levels, although occludin transcript levels were increased. Since occludin activity is highly dependent upon post-transcriptional and translational processing (Cummins, 2012), future studies will co-localize these tight junction proteins in situ to fully understand their relationship in the maternal intestine in the context of pregnancy/high fat intake.

We show that impaired maternal gut barrier integrity due to HFD intake corresponded to increased circulating maternal LPS and TNF levels. This is consistent with reports that pre-gravid obesity is associated with maternal metabolic inflammation and accumulationof CD68 macrophages inadipose tissue. We hypothesized that this low-grade systemic inflammation could alter placental function. Although the prevailing hypothesis linking maternal and offspring obesity is increased placental inflammation and macrophage infiltration (Ramsay et al., 2002; Challier et al., 2008; Zhu et al., 2010), we did not find altered pro-inflammatory cytokine transcript levels in our high-fat placentas, and found a reduction in transcript levels of a number of adaptor proteins that participate in pro-inflammatory signaling including *Traf6*, *Myd88*, *Mapk8*, and *Stat3*. As these data are limited to quantification of transcripts, evidence of placental inflammation could be present at the protein level. We did observe increased transcript levels of *Cd4*, *Cd25*, and F4/80, as well as a modest increase in F4/80 immunostaining, which is consistent with data on increased placental immune cell infiltration in humans (Challier et al., 2008; Saben et al., 2014) and in non human primates (Farley et al., 2009). These observations, combined with increased *Il10* and a reduced *Arg1*/*Nos2* ratio, suggests that placenta macrophage phenotype may be skewed by maternal HFD. Pregnancy is characterized by a state of monocyte activation and higher number of non-classical monocytes and recently, maternal pre-gravid obesity was associated with a shift towards Th2 cytokine production (Sureshchandra et al., 2018).

As alternatively activated macrophages are key regulators of placental angiogenesis (Zhao et al., 2018), a skewing of macrophage activation is consistent with the reductions we observed in placental transcript levels of key angiogenic factors. CD44 (cell surface receptor of hyaluronan) is key in tissue remodeling late in pregnancy (Marzioni et al., 2001), and bone morphogenetic proteins (BMPs) are essential for uterine vascular development, with Bmpr2 knockout mice demonstrating characteristics of preeclampsia (Nagashima et al., 2013). VIP supports extravillous trophoblast cell migration, invasion and spiral artery remodeling (Paparini et al., 2017). WNT and its receptor LRP5 signal to control cell proliferation and differentiation and a loss of WNT signaling is associated with poor placental vascular development (Ishikawa et al., 2001). Indeed, we do show evidence of increased apoptosis via increased immunostaining of AC3 in placentas from high-fat fed dams. It is possible that these signaling pathways are impaired in maternal obesity and future studies are ongoing to investigate these networks more thoroughly.

Hypoxia is a known stimulator of angiogenesis (Shweiki et al., 1992) which is consistent with our observed increase in VEGF and CD31 immunostaining. As we observe a reduction in smooth muscle actin, we hypothesize that the placentas of high-fat fed dams may have impairments blood vessel maturation, which is supported by previous reports of HFD decreasing uteroplacental blood flow (Frias et al., 2011) and vascularization (Hayes et al., 2012) in rodents, although pathways in non human primates and humans may be different (Farley et al., 2009; Lappas, 2014). Hypoxia also induces a metabolic switch from oxidative phosphorylation to glycolysis to prevent ROS production (Weckmann et al., 2018), and has previously been associated with decreased urine carnitine (Conotte et al., 2018) and placental carnitine uptake (Rytting and Audus, 2007; Chang et al., 2011). In humans, maternal HFD decreases levels of carnitine and short-and medium-chain acylcarnitines in the placenta (Calabuig-Navarro et al., 2017). We also show that maternal HFD significantly reduces placental carnitine (C0) and its acylcarnitine derivatives (C2, C3, C8, C10). Carnitine is required for mitochondrial function, and its deficiency can suppress autophagy through increased levels of acetyl-CoA (Mariño et al., 2014). Decreased placental carnitine uptake may be responsible for our observed reductions in transcript levels of autophagy-regulators *Vsp15* and *Vps34*.

Reductions in these placental metabolites could impact fetal development. Carnitine is transported across the placenta to accumulate in fetal tissues over the course of gestation (Nakano et al., 1989) and carnitine deficiency has been associated with intestinal inflammation and apoptosis in neonatal mice (Sonne et al., 2012). We found that NF-κB activity was increased in the fetal small and large intestine with maternal HFD, but this was not accompanied by increased transcription of pro-inflammatory cytokines. This may be due to insufficient p38 MAPK activation, which has previously been shown to be essential for intestinal inflammation due to NF-κB activation (Guma et al., 2011) as well as intestinal epithelial cell differentiation (Houde et al., 2001). As canonical activation of NF-κB during development reduces TNF-induced apoptosis in fetal liver and lung (Espín-Palazón and Traver, 2016), increased NF-κB may represent a homeostatic response to HFD induced increases in maternal circulating TNF levels. As amniotic fluid inhibits TLR signaling (Good et al., 2012), increased *Tlr4* transcripts in fetal intestines in association with maternal HFD may lead to increased gut inflammation postnatally. While these data do not suggest that maternal HFD intake directly increased fetal gut inflammation, they may indicate an increased susceptibility to inflammation postnatally.

TLR4 signaling through MyD88-dependent pathways has previously been shown to be required for LPS-mediated increases in intestinal tight-junction permeability (Guo et al., 2015). We observed increased mRNA levels of tight junction proteins in fetuses of high-fat fed dams, but no change in protein levels. This may be related to the observed down-regulation of the IRE1-arm of the unfolded protein response (UPR). Down-regulation of the IRE1 arm may include decreased mRNA degradation via regulated IRE1-dependent decay (RIDD), which could explain increased mRNA levels of ER-taxing proteins including *Muc2*, *Gpr43*, and occludin, as well as increased levels of *Grp78*, which is stabilized by RIDD (Kimmig et al., 2012). The importance of the UPR in the differentiation of the intestinal epithelium has previously been demonstrated (Heijmans et al., 2013) and while our data do not support the hypothesis that maternal HFD induces fetal gut ER stress and decreased barrier function, they may indicate impaired intestinal differentiation. Future work will investigate the impact of maternal HFD on fetal intestinal proliferation and differentiation using histology and immunofluorescence. These investigations were beyond the scope of the present study.

## 5 CONCLUSION

In conclusion, we show significant changes in the gut microbiota and intestinal adaptations during pregnancy in a mouse model of maternal HFD. Maternal HFD intake perturbed the gut microbiota and modified maternal intestinal adaptations to pregnancy. Maternal HFD reduced maternal intestinal barrier function, possibly by impairing tight junction integrity. Maternal HFD modestly altered the mRNA of immune cell markers in certain gut sections. Further work is required to understand the inflammatory signaling pathways that are initiated in response to altered microbiota and intestinal barrier integrity. Maternal HFD was associated with increased placental hypoxia and reduced placental carnitine-derived metabolites, as well as increased NF-κB activation and decreased activation of the UPR inthe fetal gut. Due to the significant role that gut microbiota and intestinal barrier function play in mediating HFD induced obesity, it is likely that maternal intestinal changes modulate adverse metabolic adaptations that occur during high-fat pregnancy and impart increased risk of obesity in the offspring.

## ACKNOWLEDGEMENTS

The authors would like to acknowledge the funding bodies Canadian Institutes for Health Research, Natural Sciences and Engineering Research CouncilofCanada and Genome Canada, the Farncombe Family Digestive Health Research Institute and the Canada Research Chairs Program.

## AUTHOR CONTRIBUTIONS

WG performed the animal experiments, contributed to acquisition, analysis, and interpretation of data, and contributed to writing the manuscript; KMK contributedtoacquisition, analysis, and interpretation of data and wrote the manuscript; JGW contributed to acquisition and analysis of data and edited the manuscript; MS performed metabolomics and edited the manuscript; CJB contributed to acquisition and analysis of data and edited the manuscript. PBM assisted in the metabolomics analyses and edited the manuscript; JJP acquired and analyzed CAIX, VEGF, VEGFR2, CD31, and *α*-SMA IHC/IF data and edited the manuscript; MGS assisted in the bacterial sequencing analyses and edited the manuscript; DMS designed the experiments, assisted in analyzing and interpreting the data, and prepared the manuscript. All authors have approved the final version of the manuscript and agree to be accountable for all aspects of the work. All persons listed as authors qualify for authorship, and all those who qualify are listed.

